# Another mystery snail in the Adirondacks: DNA barcoding reveals the first record of *Sinotaia* cf. *quadrata* (Caenogastropoda: Viviparidae) from North America

**DOI:** 10.1101/2021.01.28.428687

**Authors:** Ethan O’Leary, Donovan Jojo, Andrew A. David

## Abstract

Alien molluscs pose a serious threat to global freshwater diversity and have been implicated in many ecosystem-altering invasion events over the past few decades. Biomonitoring surveys are therefore a key tool for ensuring biosecurity in diversity hotspots and vulnerable habitats. In this study, we use DNA barcoding to provide the first record of the viviparid, *Sinotaia* cf. *quadrata* from North America. Reciprocal monophyly and low genetic divergence (uncorrected p-distance: 0.004) with a *Bellamya quadrata* individual from the type region (China) provides strong support for this identification. The species was recovered as part of a routine biomonitoring survey of the Adirondack region of northern New York. Only three adults were recovered (no populations or juveniles) indicating that the discovery represents a very recent arrival. Considering the proximity of the sampling site from the massive St. Lawrence River, it is likely that *S*. cf. *quadrata* was introduced into the St. Lawrence, probably via the aquarium plant trade, and was able to spread into smaller river system in northern New York and possibly other border states. This record represents the fourth alien viviparid, the third of which is of Asian origin, that have made its way to New York waters. Future biomonitoring efforts for the upcoming summer period will involve targeted searches for *S*. cf. *quadrata* to determine the extent of its spread in the region.

---

Aquatic invasive species (AIS) pose a serious threat to biological communities in both marine and freshwater systems (David and Janac 2018). In the freshwater realm, notable events such as the zebra mussel invasion of the Great Lakes of North America (Griffiths et al. 1991), the rapid spread of the New Zealand mud snail across the United States (Loo et al. 2007) and more recently, the march of the marbled crayfish across Europe (Hossain et al. 2019), has garnered significant attention both within and outside of the invasion science community. While the majority of introduced species rarely become invasive, the few that do can severely, and in some cases permanently disrupt community dynamics and ecosystem functioning, ultimately leading to biodiversity loss (Molnar et al. 2008). Early detection of AIS via frequent biomonitoring of rivers and lakes is therefore a key task that must be undertaken to help prevent future invasion events (Hamelin and Roe 2019; Pederson et al. *in press*).

The Adirondack Park, situated in northern New York, is the largest protected forest reserve in the contiguous United States, and with more than 100,000 hectares of lakes and streams, it is the largest protected freshwater system in the country (Erickson 1998). More than half of the rivers within and around the park drain into the Laurentian Great Lakes and the larger St. Lawrence River system which allows for bidirectional dispersal of organisms among these three regions (Shaker et al. 2017). During the summer months, the park is a popular destination for hikers, boaters and recreational anglers from around the country, and while New York State environmental regulators have implemented policies and rules aimed at minimizing the potential for non-indigenous species (NIS) introductions, the sheer size of the Adirondack region combined with a small workforce make enforcing such policies difficult. The lakes and rivers of the Adirondack region are now home to several non-indigenous molluscs including the zebra and quagga mussels (*Dreisenna polymorpha* and *D. bugensis*), New Zealand mud snail (*Potamopyrgus antipodarum*), and three viviparids: the Chinese and Japanese mystery snails (*Cipangopaludina chinensis* and *C. japonica*), and the banded mystery snail (*Viviparus georgianus*) (David et al. 2017; David and Cote 2019). In July 2020, three snails were collected from a boat launch site just outside the Adirondack Park and less than 1 km from the St. Lawrence River as part of a long-term biomonitoring survey for aquatic invasive species (Fig. 1). Both specimens were initially identified as ‘new’ viviparid records based on conspicuous conchological features which differed from the well-established invasive viviparids already present in the region. Here, with the aid of DNA barcoding, we report the first record of *Sinotaia* cf. *quadrata* from northern New York and as far as we know, this record is also the first report of the species within North American waterways. In addition to providing genetic confirmation of the species, we also briefly discuss possible pathways and vectors which may have been responsible for translocating it to this region.

**Fig. 1.**
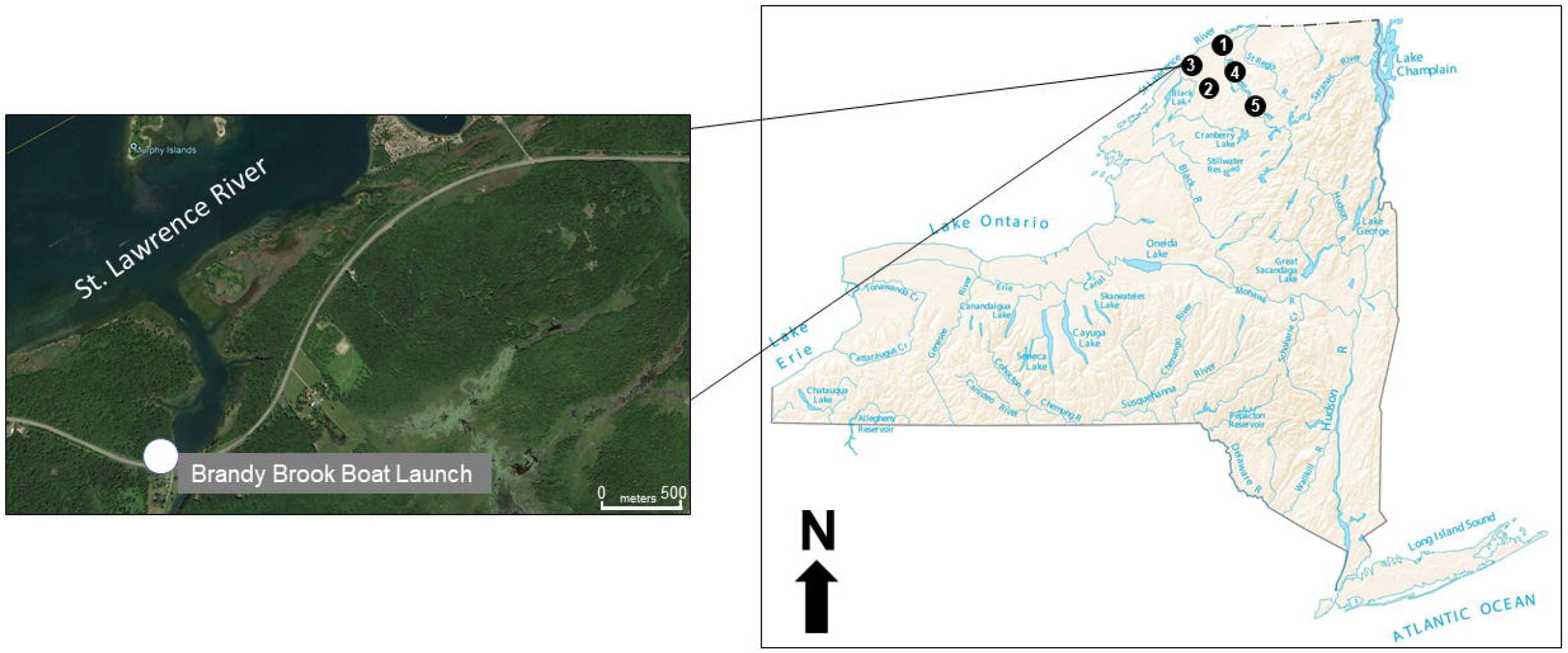
Map of New York State showing sampling localities: 1 – Massena, 2 – Heuvelton, 3 – Waddington, 4 – Potsdam, 5 – Colton. Inset map shows specific sampling site within the town of Waddington, where *Sinotaia* cf. *quadrata* was discovered.

As part of a long-term monitoring survey for AIS in the rivers and streams of the northern New York, snails were collected from five different sites in July 2020, all located at or within the vicinity of a boat launch dock (Fig. 1). Using a rapid assessment protocol outlined by David and Cote (2019), sampling was carried out at each site for exactly one hour in the shallows (maximum depth – 1.2 m). All specimens were transported live to the David Lab at Clarkson University and sorted by Family and Genus using conchological characteristics and the identification key of Jokinen (1992). Shell morphometrics were determined using the methods of Chiu et al. (2002). A Vernier caliper (error margin: 0.05 mm) was used to determine shell length (SL), shell width (SW), aperture length (AL) and aperture width (AW) and photographs of the apertural and subapertural view of each snail was taken. Forty four out of the 47 snails collected belonged to either *Cipangopaludina chinensis* or *Viviparus georgianus*, both of which are established invasive gastropods in the region. Three snails from one of the five sites (Brandy Brook Boat Launch, Waddington NY – 44°52’03.75” N, 75°11’41.90” W) exhibited a spire that was considerably shorter than the aforementioned species, though broad similarities in taxonomically informative shell traits indicated they were all viviparids This prompted a molecular investigation into their identities. All three snails were fixed in 99% ethanol for 48 hrs. for DNA barcoding. Shells of both snails were cracked and ~2 mg of tissue just above the operculum was dissected, air-dried and digested in a Proteinase K and lysis buffer solution (Qiagen, Hilden, Germany). Genomic DNA from each sample was extracted using the D’Neasy Blood and Tissue kit (Qiagen, Hilden, Germany) to produce aliquots with DNA concentrations of 85 and 149 ng/μl. A ~710 fragment of the mitochondrial gene, cytochrome *c* oxidase 1 (CO1) was amplified for both individuals using the forward and reverse primer pairs from Folmer et al. (1994): (HCO2198 and LCO 1490). Polymerase Chain Reaction was carried out in a 25 μl reaction mixture using the following cycling parameters: 92°C for 2 min, 92°C for 40 s, 51°C for 1 min, 68°C for 1 min and 68°C for 7 min. PCR products were visualized in a 1% agarose gel stained with ethidium bromide (EtBr). Amplicons were excised and cleaned using a gel purification kit (Qiagen, Hilden, Germany), and purified DNA was sequenced at GeneWiz LLC (South Plainfield, NJ, USA) using the forward primer and Big Dye Terminator Cycle Sequencing. All sequences were translated using the ExPASY online translation tool to ensure gene functionality, and then deposited into the GenBank database (accession codes: MW425606 – MW425608).

Initial molecular identification using GenBank’s BLASTn tool and the Species Level Barcode option from the Barcode of Life Database (BoLD v4) yielded a 99.09% and 99.32% similarity index with *Bellamya* (=*Sinotaia*) *quadrata*. Based on this preliminary identification, a comprehensive CO1 dataset was assembled, which included (1) representative barcodes for all known viviparid snails from New York State and (2) the sequence dataset of Arias et al. (2020), who confirmed the presence of *S. quadrata* in south-western Europe (see Table S1 in Arias et al. 2020). Sequences were aligned and edited using the Clustal W alignment tool in BioEdit ver. 7.2.5 (Hall 1999). After editing, a 435 bp fragment remained for analysis with 142 polymorphic sites, of which 124 was parsimony informative. A maximum-likelihood tree was constructed in MEGAX (Kumar et al. 2018) using the TrN + I + G nucleotide substitution model as determined by Akaike Information Criterion calculation for best fit model in jModelTest2 (Darriba et al. 2012). Pairwise uncorrected genetic distances (p-distance) were also calculated in MEGAX.

Individuals of putative *S. quadrata* individuals were dark brown with a conspicuously whitish tan on the upper whorls, which in at least one individual, extended all the way to the largest whorl (Fig. 2). Mean SL and SW were 15.4 ± 0.87 mm and 11.3 ± 0.2 mm (N=3), respectively, while mean AL and AW were 5.9 ± 0.2 mm and 3.1 ± 0.2 mm (N=3), respectively. Number of whorls ranged from 4-5 with a short, depressed spire and deep sutures especially between the largest whorls. The aperture was ovate and operculum thin with a series of slightly visible concentric growth rings. No juveniles were observed at the sampling site. Phylogenetic analysis corroborated initial BLASTn results with all three individuals nesting within a robustly supported clade which included a single *Bellamya* (=*Sinotaia*) *quadrata* individual from the type region (China) (Fig. 2). Genetic distance (p-distance) between *B. quadrata* and North America specimens was 0.004 (0.4% divergence). Genetic distances between *S. quadrata* from other localities was relatively higher (0.045 – 0.068) Two prominent clades, A and B were recovered but both had low bootstrap support: 66 and 77, respectively. Clade A consisted of species exclusively of Asian origin, including all members of the *Sinotaia* genus in the dataset along with two additional invasive viviparids in New York – *Cipangopaludina chinensis* and *C. japonica*. In contrast, Clade B was characterized exclusively by viviparids of African origin.

**Fig. 2.**
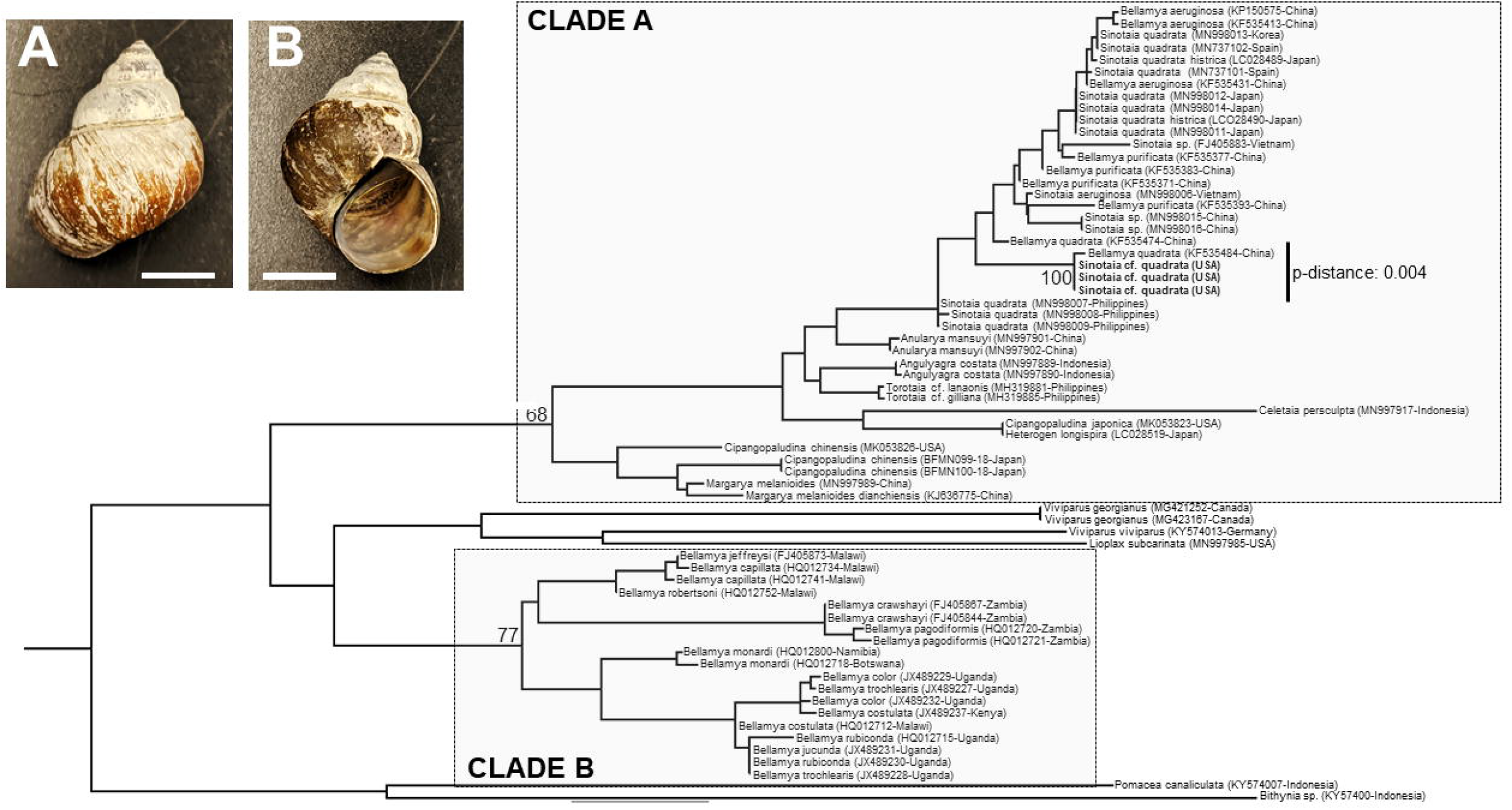
Apertural (A) and subapertural (B) photographs of *Sinotaia* cf. *quadrata* collected from northern New York (scale bars: 5 mm) along with Maximum-Likelihood phylogenetic tree of selected viviparid taxa based on 1000 bootstrap replicates; bolded taxa are those which were sequenced in this study.

Based on the results from this study, the occurrence of *Sinotaia quadrata* in New York represents the first report of the species on the North American continent and the fourth alien viviparid in New York waters, the third of which is of Asian origin. Considering that only three individuals were recovered from this survey along with the fact that the species has never been reported from the same sites in previous surveys (David et al. 2017, David and Cote 2019), their occurrence likely represents a very recent introductory event. While conchological comparisons was initially used to delineate *S. quadrata* from other viviparids in the region, these shell traits were admittedly difficult to comparatively characterize due to their variable nature among viviparids. For example, Kagawa et al. (2019) found that shell morphology of *Sinotaia* spp. in their native range can change due to environmental influences while in *Cipangopludina japonica*, significant variation in spire height and angle have been observed between specimens from its native and invasive ranges (David and Cote 2019). Like recent studies by Arias et al. (2020) and Stelbrink et al. (2020), the current study recovered two Bellamyinae clades – A and B. However, bootstrap support for both clades was lower than that reported from the Arias et al. (2020) study (68 and 77 versus 67 and 90). Phylogenetic analyses recovered a reciprocally monophyletic clade consisting of North American specimens and a single specimen (*Bellamya quadrata*) from China. Furthermore, the Chinese *B. quadrata* along and specimens from New York exhibited low genetic divergence (0.4%). Interestingly, genetic divergence across all *S. quadrata* specimens was highly variable; as low as 0.4% and as high as 7%. Such a wide range of genetic distances within a supposedly singular taxa is concerning because it is indicative of a species complex. Species complexes are usually the result of poor taxonomy. Indeed, Stelbrink et al. (2020) suggested that the taxonomic landscape of river snails in general is poorly understood, largely due to inadequate sampling resulting in incomplete phylogenetic datasets. Nevertheless, phylogenetic based identification and low genetic divergence in the COI gene has unequivocally confirmed the presence of a *Sinotaia* species for the first time in North America. Considering the problematic taxonomy of *Sinotaia quadrata*, we herein refer to the three individuals as *Sinotaia* cf. *quadrata* instead of using the nominal name, on condition that future examination of the species using a more comprehensive dataset (specifically one that involves additional genetic markers and larger sample sizes) will be needed before any definitive delineation occurs.

*Sinotaia quadrata* (Benson, 1842) is native to Asia, specifically China, Taiwan and North Korea and has been introduced to the neighboring territories of Japan, Thailand and the Philippines. Outside of Asia, the species has also been reported from South America (Ovando and Cuezzo 2012) and Europe (Cianfanielli et al. 2017; Arias et al. 2020). Its introduction to Italy was attributed to the food trade where batches of snails could have been imported by immigrant Asian communities for food markets (Cianfanelli et al. 2017) while in Spain, the aquarium plant trade has been suggested as a more likely possibility (Arias et al. 2020). The New York site where the three individuals were recovered is a popular boat launch dock site located less than a kilometer south of the St. Lawrence River (Fig. 1). A study by Cohen et al. (2007) found that the aquarium plant trade in the city of Montreal has been responsible for the introduction of thousands of alien plant propagules into the St. Lawrence Seaway – a series of canals that connect the Great Lakes to the Atlantic Ocean. Consequently, it is very likely that *S*. cf. *quadrata* could have hitchhiked on one or more of these alien plants and once it was released into the St. Lawrence River, was able to disperse into the northern New York river system. The presence of only three individuals without any juveniles indicate that a population has yet to become established in the region. If *S. cf. quadrata* does spread deeper into the Adirondack Park, its invasive potential will be high. The high fecundity and brooding behavior of *S. quadrata* (the latter resulting in increased survivorship rates) combined with the fact that viviparids have been found to exhibit a broad tolerance to various environmental stressors including as pH and temperature (David and Cote 2019; David et al. 2020) means that the species can very well become the third invasive viviparid from Asia to become established in the Adirondack Park. Ovando and Cuezzo (2012) found that *S. quadrata* in South America can alter native plant biomass which in turn can cause disruptions in community structure. Like *Viviparus georgianus, S. quadrata* has also been observed to be an egg predator of freshwater fishes, specifically bluegill fish (Nakao et al. 2006). In the Adirondack region, a 32 km lake known as ‘Black Lake’ harbors a sustainable population of bluegills and is frequented by anglers during the summer months. This lake merges with the Oswegatchie River, which flows for eight kilometers before emptying into the St. Lawrence River. If *S. quadrata* were to spread into this lake basin, it may pose a serious threat to bluegill fishes.

Currently, we are in the process of alerting the New York State Department of Environmental Conservations (NYS-DEC) to this new record, in addition to preparing materials such as placards and pamphlets to local anglers and recreational boaters for the upcoming summer season. Furthermore, we plan to intensify biomonitoring efforts at additional points of entry into the Adirondacks in early spring and throughout the summer to determine whether additional individuals or a population has become established.

## ACKNOWLEDGEMENTS

Funding for this project was provided by the New York Power Authority’s St. Lawrence River Research and Education Fund (SLRREF).

